# A privileged ER compartment for post-translational heteromeric assembly of an ion channel

**DOI:** 10.1101/2025.01.30.635714

**Authors:** Sudharsan Kannan, William Kasberg, Liliana R. Ernandez, Anjon Audhya, Gail A. Robertson

## Abstract

Mechanisms underlying heterotypic subunit assembly of ion channels and other oligomeric complexes are poorly understood. In the human heart, heteromeric assembly of two isoforms encoded by the *human ether-à-go-go related gene* (*hERG*) is essential for the normal function of cardiac I_Kr_ in ventricular repolarization, with loss of hERG1b contributing to arrhythmias associated with long QT-syndrome (LQTS). While hERG1a homomers traffic efficiently to the plasma membrane, hERG1b homomers are retained in the endoplasmic reticulum (ER). When expressed together, the two subunits avidly associate during biogenesis. Seeking rules specifying heteromeric association, we characterized the fate of hERG1b proteins using confocal and superresolution imaging in fixed and live HeLa cells. We found hERG1b sequestered in punctate intracellular structures when expressed alone in HeLa cells. These puncta, which depend on the presence of an N-terminal “RXR” ER retention signal, represent a privileged ER sub-compartment distinct from that containing ER-retained, type 2 (hERG-based) LQTS mutant proteins, which were rapidly degraded by the proteasome. Introducing hERG1a to cells with preformed hERG1b puncta dissolved these puncta by rescuing extant hERG1b. Rescue occurred by association of fully translated hERG1b with 1a, a surprising finding given previous studies demonstrating cotranslational heteromeric association. We propose that sequestration limits potentially deleterious surface expression of hERG1b homomeric channels while preserving hERG1b for an alternative mode of heteromeric hERG1a/1b channel assembly post-translationally. These findings reveal a surprising versatility of biosynthetic pathways promoting heteromeric assembly.

**Significance Statement:** hERG potassium channels are essential for repolarizing the ventricular action potential. Heteromeric hERG channels, which conduct cardiac I_Kr_, are formed by association of hERG1a and 1b subunits. However, mechanisms governing the assembly of hERG and other heteromeric protein complexes are poorly understood. We identify a noncanonical ER compartment that sequesters hERG1b, distinct from previously described ER quality control pathways that degrade misfolded proteins, including LQTS-associated hERG variants. Instead, this compartment preserves hERG1b and facilitates its post-translational assembly with hERG1a, challenging the current view of simultaneous, cotranslational heteromeric assembly. Our findings redefine ER retention as an active regulatory step in protein complex formation that may broadly govern hetero-oligomeric assembly in many systems and facilitate therapeutic approaches to arrhythmia and other diseases.

## Introduction

The *human ether-à-go-go-related gene* (*hERG*, or *KCNH2*) encodes the voltage-gated potassium channel Kv11.1, critical for repolarizing the ventricular action potential (1, 2). Loss of hERG function due to unintended channel block or genetic mutations in *KCNH2* can prolong the ventricular action potential, leading to drug-induced or type 2 Long-QT syndrome (LQT2) and sudden cardiac death (1-3). In addition to cardiac dysfunction, central nervous system diseases such as epilepsy and schizophrenia can also arise from mutations and mis-splicing of hERG (4, 5). The majority of LQT2 mutants and the schizophrenia variant of hERG are trafficking-deficient (6). Thus, understanding the mechanisms that regulate the expression, assembly, and trafficking of hERG channels is of broad biological and medical significance.

In human ventricular cardiomyocytes, alternate transcripts encode hERG1a and 1b subunits (or ERG1a and 1b in other organisms), which assemble to form heterotetrameric channels producing the repolarizing I_Kr_ current (7-12). The subunits are identical except in their N-termini, which in ERG1a harbors a Per-Arnt-Sim (PAS) domain (13) that is missing in ERG1b (7). In the mature channel, the PAS domain interacts with the C-terminus of the adjacent subunit (14), and perturbation of this interaction alters channel function (15). Expressed independently in heterologous systems, ERG1a subunits form homomeric channels that efficiently traffic to the plasma membrane and produce potassium currents. In contrast, ERG1b homomers traffic inefficiently, producing little or no current (7, 16). When expressed together, ERG1b markedly alters the gating properties of ERG1a currents, providing strong evidence for their coassembly (7). This association appears to occur cotranslationally, based on the following evidence: 1) isolated hERG1a and 1b N-terminal fragments interact in vitro in a dose-dependent manner (17); 2) coexpression of a hERG1a N-terminal fragment disrupts channel oligomerization in cells and impairs core glycosylation, a process known to occur cotranslationally (17, 18); and 3) immunoprecipitation from HEK293 cell or cardiomyocyte lysates using a hERG1a-specific antibody copurifies nascent hERG1a and 1b subunits along with their encoding mRNAs (19). Together these findings indicate that hERG1a/1b subunit assembly begins cotranslationally, early in biogenesis.

Several lines of evidence underscore the critical role of the heteromeric composition of cardiac I_Kr_. In canine adult cardiomyocytes, specific antibody staining reveals an approximately equal distribution of the two subunits, localized primarily to the Z line (9). Similarly, in both canine and human myocardium, western blot analysis using a pan-hERG antibody detects roughly equal amounts of the proteins, supporting the conclusion that cardiac I_Kr_ is predominantly formed by heteromeric hERG1a/1b channels (9). The native current also displays properties more consistent with the heteromer than the hERG1a homomer (20). The hERG1b subunit modulates hERG1a properties, yielding larger currents by accelerating the time course of activation and recovery from inactivation, thus enhancing repolarization (10). Computational studies incorporating these differences in channel gating behavior into a cardiomyocyte model predicted that loss of hERG1b would be pro-arrhythmic (10). This prediction was later validated experimentally: shRNA depletion of hERG1b in cardiomyocytes or introduction of the hERG1a-specific N-terminal PAS domain in trans, which effectively converts native heteromers into hERG1a homomers, leads to prolonged action potential duration (APD) and increased APD variability, two cellular markers of pro-arrhythmia (8).

Previous studies identified an N-terminal, arginine-based endoplasmic reticulum (ER) retention motif (RXR) in hERG1b that is critical for its retention in the ER in the absence of hERG1a (16). In heterologous expression systems, mutating the RXR motif effectively rescues hERG1b from ER retention: hERG1b proteins are fully glycosylated (mature) as assayed by western blot, and they produce robust currents, indicating successful plasma membrane expression (16). Thus, RXR-mediated ER retention ensures that hERG1b subunits predominantly reach the plasma membrane as components of heteromers, while restricting their expression as homomers. However, the detailed mechanisms of ER retention and rescue of 1b remain unclear.

In this study, we discovered that retained hERG1b proteins are stably sequestered within a privileged ER compartment, which we have termed the “ER sequestration and assembly” (ERSA) compartment. This compartment provides a protected environment, shielding hERG1b from degradation while it awaits assembly with hERG1a, adding a new level of complexity to the previously reported mechanism of cotranslational assembly (17, 19). These findings reveal that biogenesis of heteromeric I_Kr_ channels is prioritized through two distinct mechanisms.

## Results

### hERG1b is sequestered in intracellular puncta

hERG1a and 1b isoforms exhibit distinct cellular distributions when expressed heterologously. Immunofluorescence confocal microscopy in HeLa cells shows that hERG1a displays a widespread ER signal along with discernible plasma membrane localization, indicative of mature, functional membrane expression (Fig. 1*A*) (21). In contrast, hERG1b forms numerous large intracellular puncta with fluorescence signals measuring 300 nm or greater, detected using automated image analysis (Fig. 1 *B* and *C*; see Methods). Many fewer puncta are observed for hERG1a (Fig. 1 *A* and *C*). A corresponding difference is observed when the two isoforms are expressed exogenously at low levels in the more native-like milieu of live cardiomyocytes derived from iPSC cells (Fig. 1 *D-F*).

**Fig. 1.**
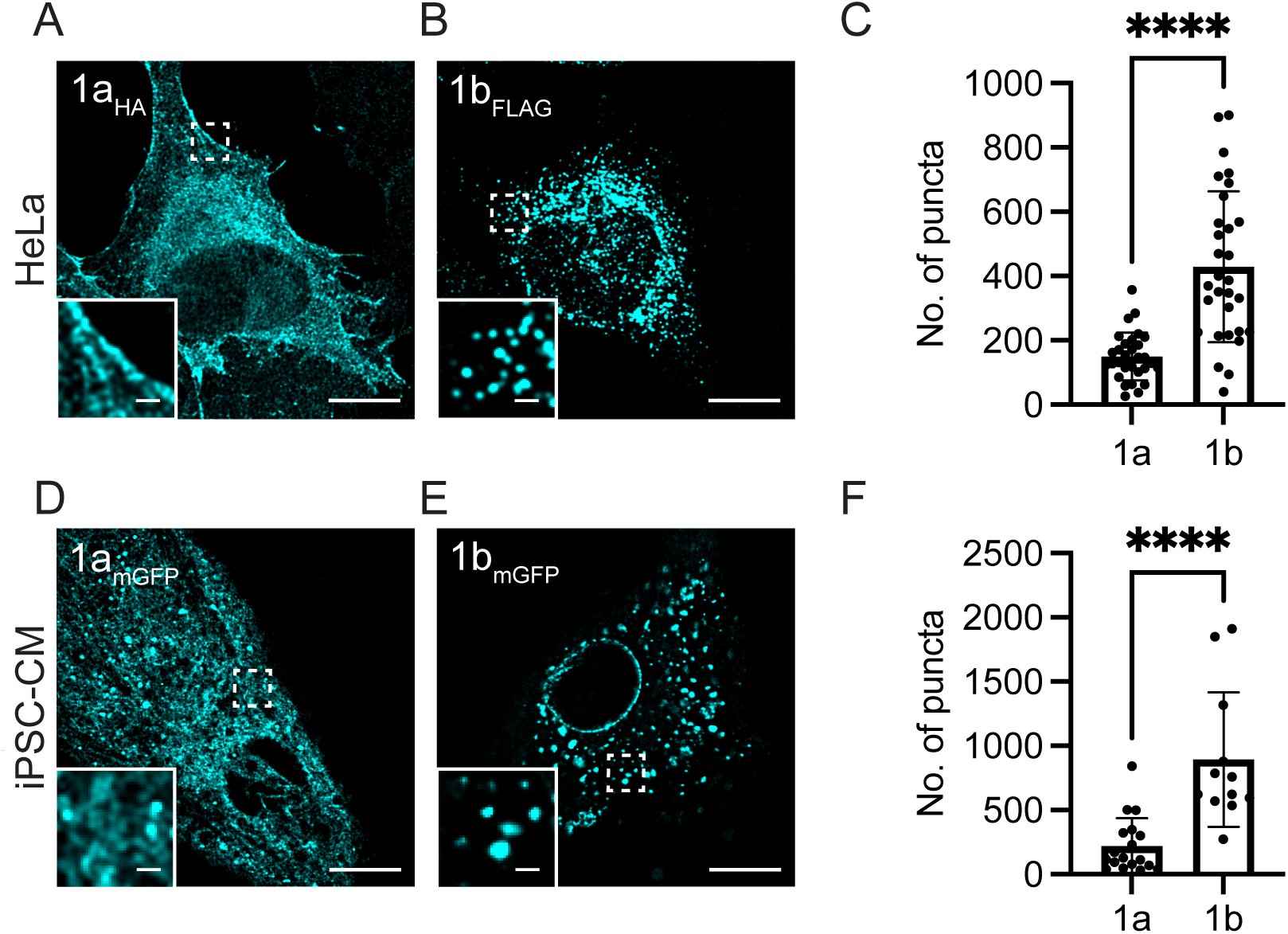
hERG1b is sequestered in intracellular puncta. (*A*) Confocal images of fixed and immunostained HeLa cells transfected with *hERG1a* or (B) *hERG1b* plasmids; (*C*) quantification of the number of puncta per cell; data are mean ± s.d., n= 36 cells per condition, analyzed with Student’s t-test (**** indicating p<0.0001); (*D*) confocal live cell images of iPSC-CM transfected with 1a-GFP or (E) 1b-GFP; (*F*) quantification of number of puncta per cell; data are mean ± s.d., n=18 cells for 1a_GFP_, 12 cells for 1b_GFP_, analyzed with Student’s t-test (**** indicating p<0.0001); Scale bar: 10 μm in large image and 1 μm in the inset.

Control experiments confirm that these puncta are not an artifact of the C-terminal tags used for detection. Untagged hERG1b puncta expressed in HeLa cells showed the same distribution when stained with pan-hERG antibody (Fig. S1*A*), and native puncta were also observed in iPSC-CMs using a pan-hERG antibody (Fig. S1 *B* and *C*), although the lack of availability of a hERG1b-specific antibody precluded isoform assignment. Notably, a previous study from our lab demonstrated a punctate localization pattern for hERG1b in antibody-stained canine ventricular myocytes, whereas hERG1a exhibited a more diffuse distribution (9). These findings suggest that the puncta serve as the intracellular substrate for the well-documented ER retention of homomeric hERG1b subunits (10, 16, 17).

### hERG1b puncta are localized to ER

To determine the subcellular localization of the puncta at superresolution, we imaged hERG1b in iPSC-CMs using two-color stimulated emission depletion (STED) microscopy (Fig. 2). ER labeling was achieved by fusing HaloTag to an ER signal sequence and the KDEL ER retention motif (Halo-KDEL) (22), while hERG1b was tagged with a C-terminal SNAP tag (1b_SNAP_). mRNA was transcribed in vitro from the Halo-KDEL and 1b_SNAP_ plasmid constructs and transfected into iPSC-CMs. After 24h, fluorescent labeling of the ER with the JFX650 HaloTag ligand and 1b_SNAP_ with the JFX554 SNAP-tag ligand revealed that more than 95% of the puncta were centered within a 50 nm distance from the ER membrane, confirming their ER localization (Fig. 2 *A-E*).

**Fig. 2.**
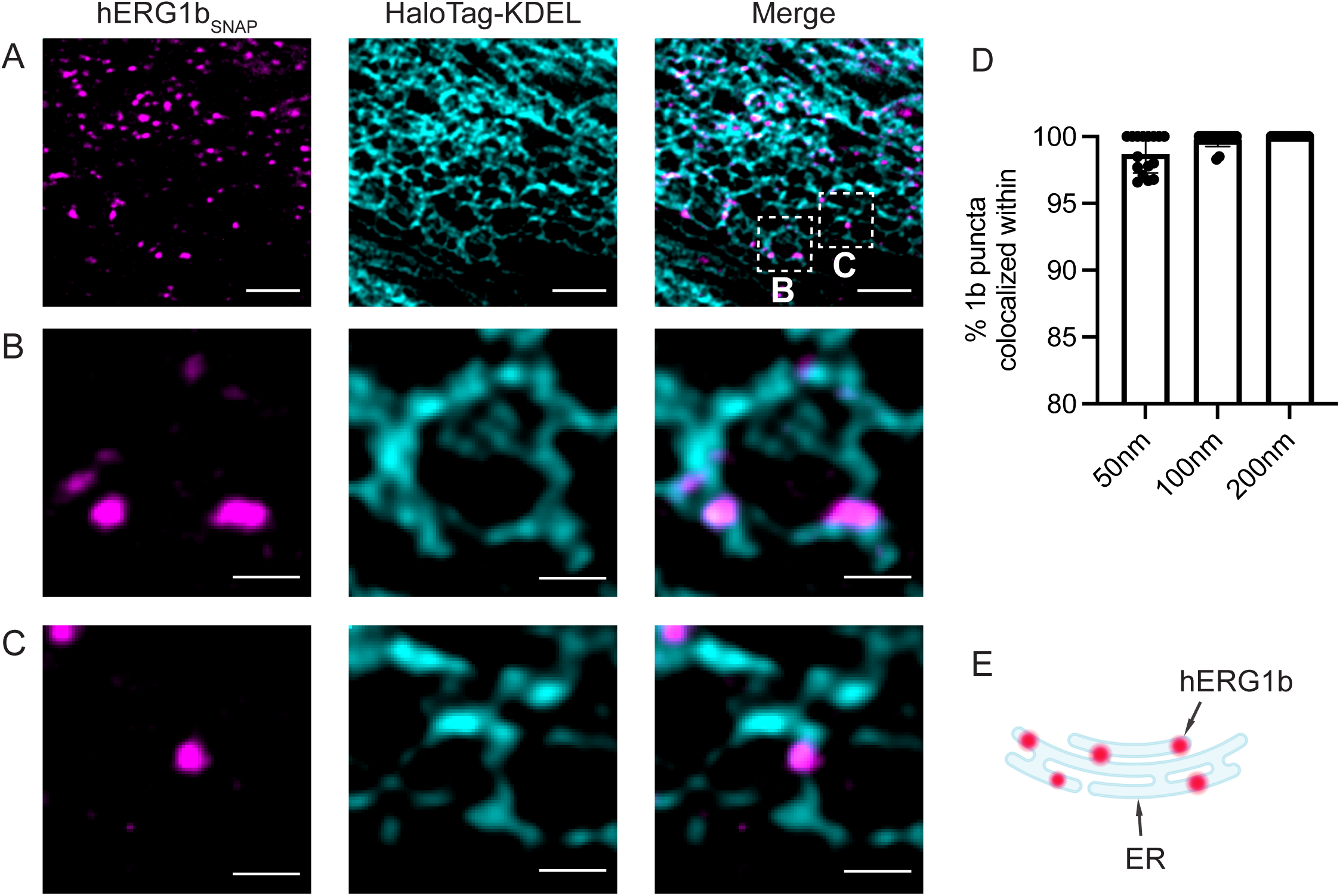
hERG1b puncta are localized to ER. (*A-C*) STED microscopy of iPSC-CM transfected with mRNAs encoding hERG1b_SNAP_ labeled with JFX554 (magenta) and ER marker Halo-KDEL labeled with JFX650 (cyan); (*D*) quantification of colocalization events thresholded with three different distances between the center of hERG1b puncta and the surface of the ER; data are mean ± s.d., n= 16 regions of interest (see Methods); (*E*) cartoon of ER localization of hERG1b puncta. The scale bar is 10 μm for *(A)* and 500 nm for (*B,C*).

hERG1b puncta showed minimal colocalization with other subcellular compartments, including COPII (ER exit), COPI (Golgi-to-ER retrieval), lysosomes, or autophagosomes (Fig. S2). Interestingly, the presence of few autophagosomes (Fig. S2*C*) indicates that hERG1b expression did not trigger significant cellular stress under these conditions (23). These findings establish that hERG1b puncta represent an ER-resident substructure.

### hERG1b is sequestered in a privileged ER compartment

A majority of LQT2-mutant hERG channels fail to mature, accumulating in the ER where they are ubiquitinated and targeted for proteasomal degradation (6, 24-27). To determine whether ER-sequestered hERG1b meets a similar fate, we compared its retention to that of the trafficking-deficient LQT2 mutant hERG1a-Y611H using HA-tagged constructs transfected into HeLa cells and immuno-stained (25). As expected, hERG1a-Y611H exhibited intracellular signal with no apparent membrane localization, consistent with its ER retention phenotype; however, it exhibited relatively few puncta compared with hERG1b (Fig. 3 *A-C*).

**Fig. 3.**
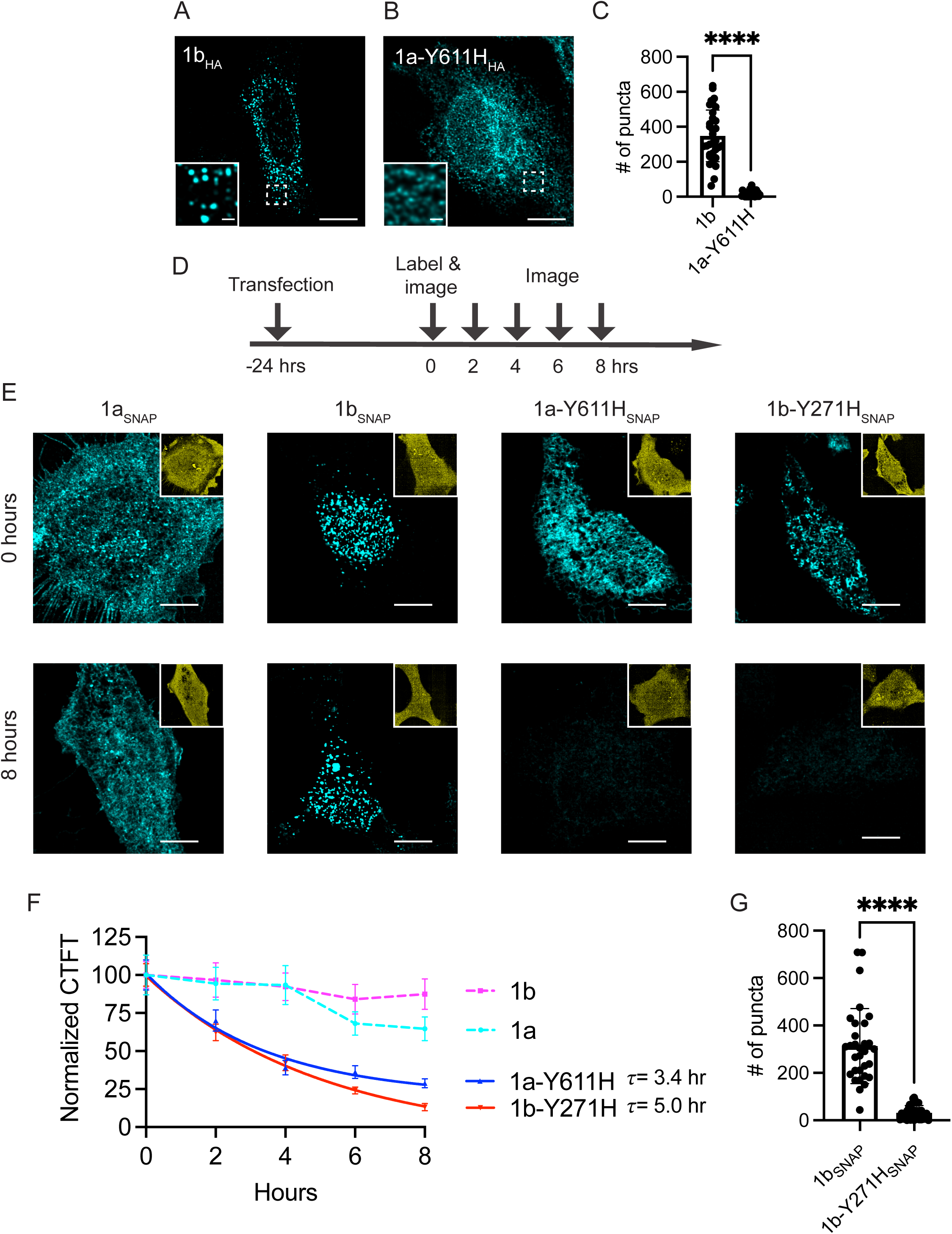
hERG1b is sequestered in a privileged compartment. (*A*) Immunostaining of HeLa cells transfected with hERG1b_HA_ or (*B*) hERG1a-Y611H_HA_; (*C*) quantification of the number of puncta per cell; data are mean ± s.d., n= 33 for hERG1b, n= 26 for 1a-Y611H, analyzed with a Student t-test (**** indicating p<0.0001); (*D*) experimental design for the pulse-chase experiment; (*E*) HeLa cells transfected with either hERG1a_SNAP_, 1b_SNAP_, 1a-Y611H_SNAP_ or 1b-Y271H_SNAP_ imaged at various time points provided in (*D*) and sample images provided for 0 hr and 8 hr after labeling; (*F*) quantification of normalized corrected total fluorescence intensity (CTCF) plotted across time; hERG1a-Y611H and hERG1b-Y611H data are fitted with single exponential decay; hERG1a, hERG1b data are plotted with connected lines; data are mean ± SEM, n= 35-45 cells per condition; data with individual data points are provided in Fig. S4; (G) quantification of the number of puncta per cell; data are mean ± s.d., n=31 for hERG1b, n=32 for 1a-Y611H, analyzed with a Student t-test (**** indicating p<0.0001); inset on the upper right of (*E*) shows the Halo-tag expression (yellow) used to identify the transfected cells. Scale bar: 10 μm for large images (*A-E*) and 1 μm for insets (*A, B*).

To determine relative stabilities, we designed a pulse-chase paradigm to monitor degradation of SNAP-tagged constructs labeled with fluorescent ligand (Fig. 3 *D-F*). Cells were identified with HaloTag expressed independently from an internal ribosome entry site (IRES) in the plasmid encoding each isoform (upper right insets, yellow). After 24 h transfection, we labeled extant SNAP-tagged proteins with fluorescent ligand JFX650 (cyan) and then blocked detection of all subsequently synthesized protein using a non-fluorescent ligand (PA-JF-549); thus, proteins labeled and imaged at t=0 could be imaged again at 2-h intervals after labeling (Fig. 3 *D*). As previously demonstrated, hERG1a fluorescence exhibited only a modest reduction over the 8-h period whereas hERG1a-Y611H fluorescence showed a relatively rapid, single-exponential decay, presumably via the action of the proteasome on ER-retained protein (25). The corresponding LQT2 mutation in hERG1b showed a similar decay. In contrast, hERG1b exhibited the greatest stability over time (Fig. 3 *E-F*; Fig. S3). These findings indicate that wild-type hERG1b is harbored in the ER in a privileged compartment we have termed “ER sequestration and assembly compartment,” or ERSA. These experiments show that ERSA is distinct from ER-associated compartments mediating proteolysis and cell quality control in a disease setting. Notably, introducing Y271H mutation reduced the number of hERG1b puncta and led to a significant reduction in protein levels in 8 h (Fig 3. *E-G*), suggesting hERG1b ERSA precludes misfolded proteins.

### An ER retention motif promotes sequestration of hERG1b in ERSA

Previous studies showed that hERG1b trafficking is promoted when an N-terminal ER retention motif, R^15^XR, is mutated. Enhanced trafficking is reflected in the maturation of a lower, core-glycosylated band on a western blot to a higher, fully-glycosylated band, and by the expression of membrane currents (16). We determined the effect of this mutation, N^15^XN, on the punctal content of cells, finding the number reduced by about half (Fig. 4 *A*, *B*, and *D*). A quantitatively similar result was observed upon coexpression of hERG1b with hERG1a, known to “rescue” hERG1b in maturation assays (16, 28).

**Fig. 4.**
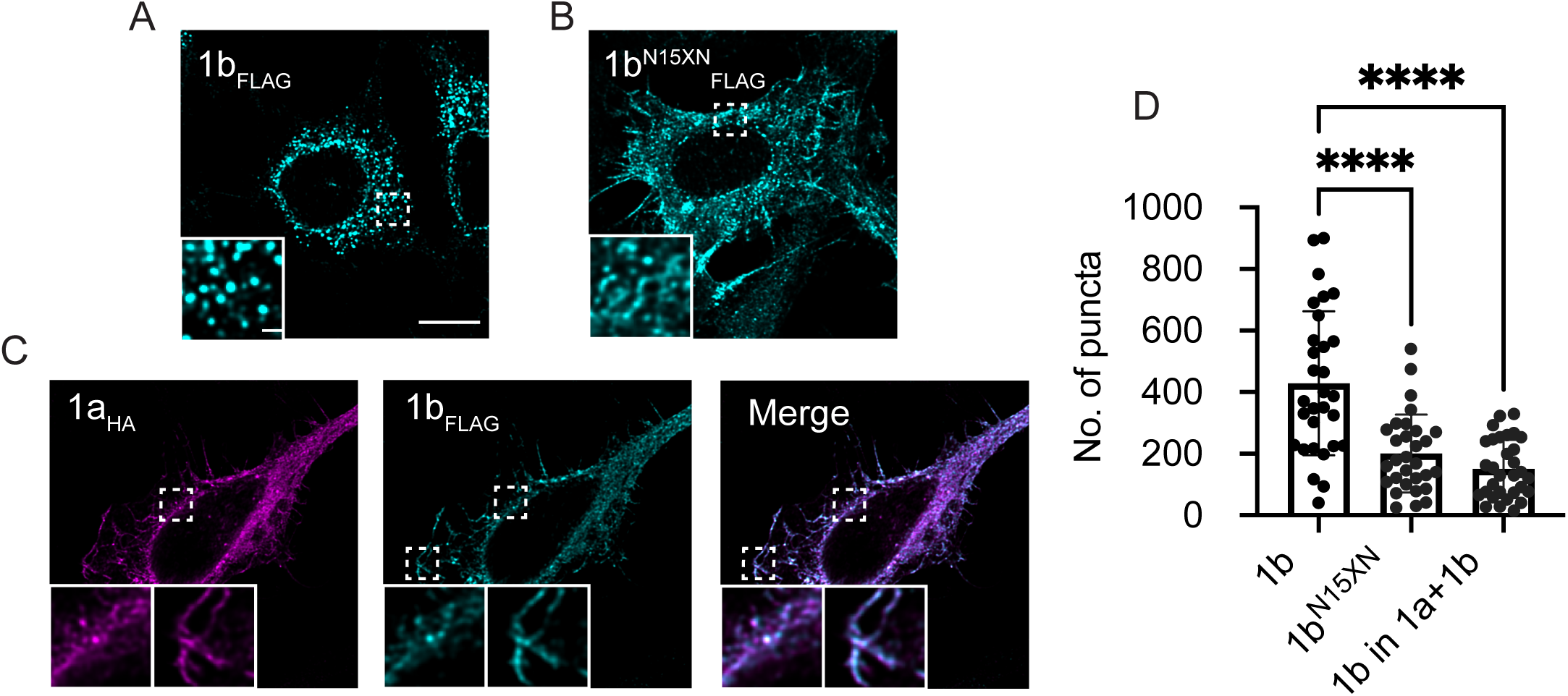
An ER retention motif promotes sequestration of hERG1b in ERSA. (*A*) confocal images of HeLa cells expressing FLAG-tagged hERG1b alone, (*B*) hERG1b with N^15^XN mutation, and (*C*) co-expression of WT hERG1a and hERG1b; (*D*) quantification of the number of puncta per cell; data are mean ± s.d. (n= 30 cells per condition) analyzed with a one-way ANOVA (**** indicating p<0.0001); scale bar: 10 μm in large image and 1 μm in inset.

Indeed, we found that coexpression with hERG1a yields fewer hERG1b puncta and the signal was instead distributed throughout the cytoplasm and plasma membrane (Fig. 4 *C* and *D*). Thus, association with hERG1a dissolves the 1b puncta or prevents their formation. Moreover, these findings establish the puncta as the physical substrate of previously described hERG1b ER retention.

### Are hERG1b puncta condensates?

We hypothesized that hERG1b puncta could represent either membranous vesicles or condensates arising from multivalent interactions of the cytosolic channel domains (29). To test whether hERG1b puncta are enclosed by membrane, we labeled cellular lipids with Potomac Yellow dye (30, 31). Amid the diffuse staining of organelles, we observed puncta that we assume represent membranous vesicles (Fig. 5 *A*, left panel). However, we observed only limited colocalization with fewer than 5% of the Potomac Yellow spots overlapping with hERG1b puncta using a 300 nm center-to-center distance threshold (Fig. 5 *A* and *B*). We conclude that hERG1b is not bound in membranous vesicles. Instead, hERG1b proteins concentrate in puncta that we operationally term condensates.

**Fig. 5.**
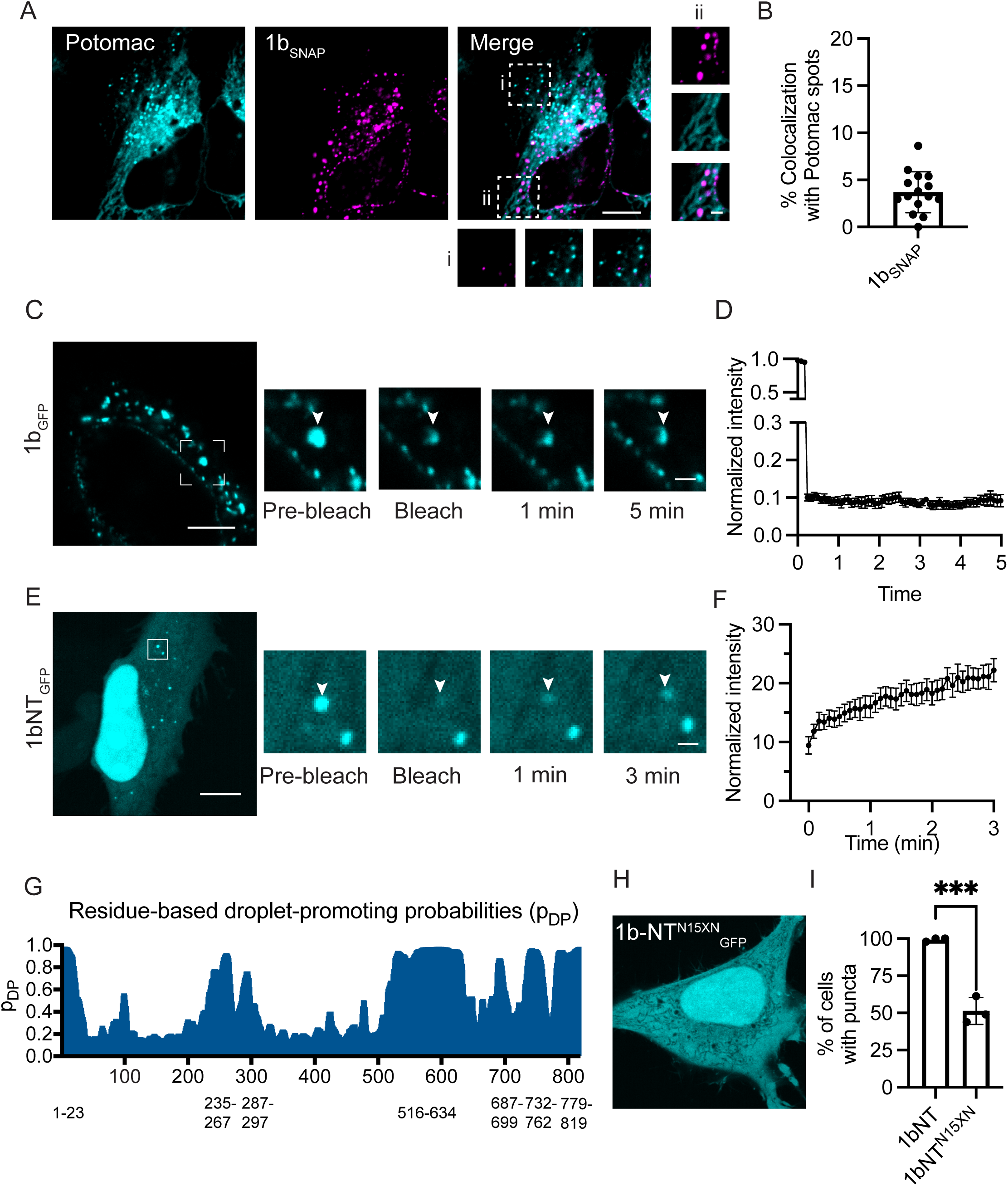
Are hERG1b puncta are condensates? (*A*) Confocal images of live HeLa cells expressing hERG1b (magenta) stained with Potomac-Yellow dye (cyan) labeling membranes; scale bar: 10 μm in the large image and 2 μm in the insets (i, ii); (*B*) quantification of percentage of number of hERG1b puncta colocalizing within 250 nm with Potomac-labeled membranous puncta (n=15 cells); (*C*) confocal images of live HeLa cell expressing hERG1b_GFP_ with photobleaching of individual puncta imaged before and after photobleaching; scale bar: 5 µm in large image and 1 µm in insets; (*D*) quantification of fluorescence intensity of the photobleached puncta before and after photobleaching; data are mean ± SEM; n=10 puncta from different cells. Data are from a representative experiment repeated three times; (*E*) GFP tagged hERG1b amino-terminal domain (1-64 residues) undergoes phase separation, with fluorescence recovery after photobleaching of individual puncta; (*F*) quantification of normalized fluorescence intensity after photobleaching; data are mean ± SEM; n= 13 puncta from different cells in a representative experiment repeated three times; (*G*) droplet-promoting regions in hERG1b identified by Fuzdrop, which includes the N-terminal domain. (*H*) Confocal image of HeLa cell expressing 1bNT^N15XN^_GFP_; scale bar: 10 μm in large image and 1 μm in insets; (*I*) graph showing % cells with puncta in 1bNT-WT and 1bNT-N_15_XN mutant protein-expressing cells; 1bNT-WT exemplar is shown in *E*; n= 3 independent wells per condition, with an average of 50 cells in each well, data are mean ± s.d., analyzed with a Student’s t-test (*** indicating p<0.001).

If ERSA condensates arise from liquid-liquid phase separation, we would expect to observe dynamic movement of molecules, which can be assessed using fluorescence recovery after photobleaching (FRAP). With this technique, recovery of fluorescence would reveal the diffusion of fluorescently tagged protein into the bleached area (32). However, we observed no FRAP of puncta for at least 5 min after photobleaching (Fig. 5 *C* and *D*). In addition, the puncta were insensitive to treatment with 1,6-hexanediol, a treatment often used to disrupt weak hydrophobic interactions promoting phase separation (33) (Fig. S4). Thus, hERG1b condensates may have non-diffusive, gel-like properties like ER-based TIS granule mesh-like condensates (34, 35), or arise as a consequence of association within a lipid raft or via protein scaffolding (36, 37).

Given that the differences between hERG1a and 1b occur solely within their N-terminal domains, we assayed the behavior of GFP-tagged hERG1b N-terminal fragments (1bNT). We found fluorescent signals in scattered cytosolic puncta and densely concentrated in the nucleus (Fig. 5*E*). The cytosolic puncta were dynamic, exhibiting FRAP with slow recovery (Fig. 5 *E* and *F*). In silico analysis of hERG1b via Fuzdrop, a proteome-wide ranking of proteins according to their propensity to form biomolecular condensates (38), reveals likely droplet-forming regions in several positions of hERG1b including the 1b-specific N-terminus (Fig. 5*G*). Thus, the hERG1b N-termini have characteristics that facilitate their dynamic homotypic association. Like the full-length protein, mutating R^15^XR to N^15^XN reduced the propensity of 1bNT to form puncta, as quantified by counting the percentage of cells with puncta (Fig. 5 *H* *and I*), mirroring its effect on full-length hERG1b and highlighting a key role for this motif in the condensate. Whether the N-terminal propensity to phase separate plays a role in hERG1b condensation in puncta will require further investigation.

### Sequestered hERG1b is rescued by hERG1a post-translationally

In previous studies we showed that hERG1a and 1b assemble cotranslationally as the subunits emerge from the ribosome (17, 19). Based on this model, we predicted that hERG1a rescues hERG1b cotranslationally by associating with newly synthesized subunits, while the sequestered hERG1b would be degraded via quality control mechanisms. However, our current findings that hERG1b is stably retained in the ERSA compartment prompted us to test whether pre-existing hERG1b proteins could be rescued by hERG1a.

To accomplish this experiment, we created a stable cell line inducibly expressing hERG1b_SNAP_ and tracked its fate in live cells. After inducing hERG1b_SNAP_ for 18 h, we labeled the existing hERG1b proteins using a 30-min pulse of fluorescent JFX650 SNAP-tag ligand. We then added a nonfluorescent PA-JF-549 SNAP-tag ligand to the media to block labeling of newly synthesized proteins (Fig. 6*A*). Cells were then transfected with hERG1a-GFP. After 24 h, we observed few hERG1b puncta and clear membrane localization of both hERG1a and 1b compared to mock-transfected controls (Fig. 6 *B-D*), indicating that, unexpectedly, existing hERG1b subunits were rescued by hERG1a. Thus, the ERSA both protects proteins from degradation and ensures they are readily available for association with newly synthesized hERG1a.

**Fig. 6.**
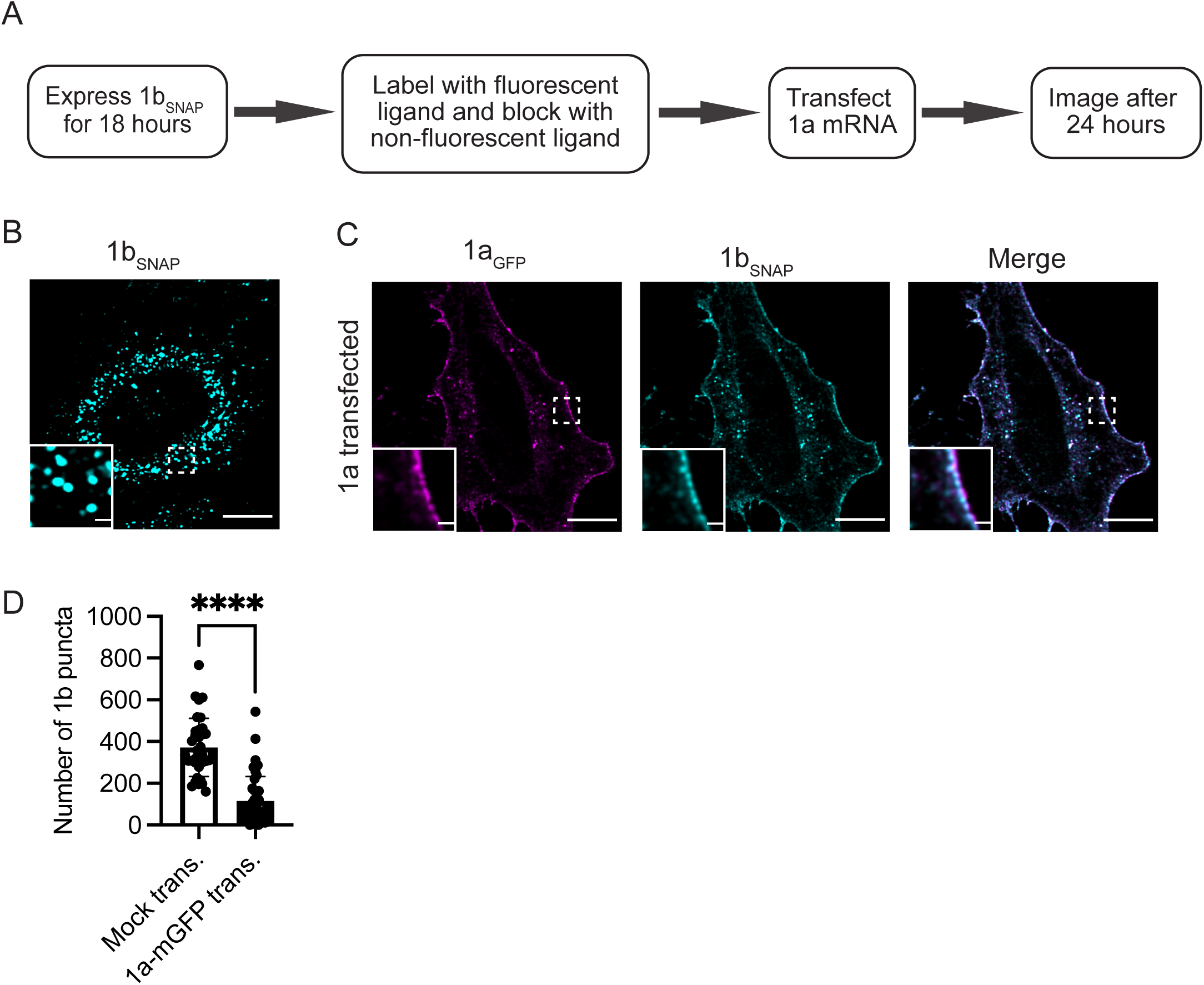
Sequestered hERG1b proteins can be rescued by newly expressed 1a. (*A*) Experimental design; (*B*) mock-transfected cells showing hERG1b puncta; (*C*) 1a-transfected cells showing hERG1b signal in or near plasma membrane in live cell imaging; (*D*) quantification of the number of puncta per cell. Data are mean ± s.d., n= 35-42 cells per condition, and analyzed with a Student’s t-test (**** indicating p<0.0001). Scale bar: 10 μm in large image and 1 μm in inset.

## Discussion

In this study, we demonstrate that hERG1b proteins, in the absence of hERG1a, are selectively retained in a punctate ER sequestration and assembly compartment, or *ERSA*. ERSA shields wild-type hERG1b proteins from proteasomal degradation, unlike LQT2 mutant hERG1a and 1b proteins, which are rapidly degraded. Mutating an N-terminal ER retention signal or co-expressing with hERG1a reduces hERG1b occupancy within ERSA, indicating that this compartment serves as the physical substrate for retained hERG1b. Notably, hERG1a rescues fully synthesized hERG1b, highlighting a novel role for ERSA in facilitating hERG channel heteromerization beyond previous models of coordinate cotranslation (17, 19). ER sequestration thus acts as a protective mechanism, safeguarding hERG1b from proteasomal degradation and preserving it in space and time until hERG1a becomes available for heteromeric assembly.

Recent studies describe macromolecular protein assembly via three pathways: (1) “co-cotranslational” or simultaneous assembly, where nascent proteins interact as they emerge from the ribosome; (2) “co-post-translational” or sequential assembly, where a fully formed subunit assembles with a different, nascent protein; and (3) “post-translational” assembly where fully formed subunits assemble after translation is complete (39, 40). Previous evidence supports the idea that hERG1a and 1b assemble through simultaneous cotranslation: in vitro, their heterotypic N-terminal domains associate in a dose-dependent manner; and in cells, disrupting N-terminal interactions disrupts oligomerization and concomitantly diminishes core glycosylation, which serves as a time stamp for early biogenesis (17, 18). Here, we present a new paradigm in which fully synthesized hERG1b can also associate with nascent or newly synthesized 1a, as demonstrated for soluble proteins assembling to form macromolecular structures in yeast (41, 42). It is unclear whether this association follows a co-post-translational mechanism where 1a is translated near hERG1b, or a strictly post-translational mechanism, in which the hERG1b-containing ERSA compartment is dispersed when in contact with full-length hERG1a subunits.

Findings from this study raise the question of stoichiometry of hERG1b subunits within the ERSA compartment. One possibility is that hERG1b subunits assemble into homotetramers, which are escorted out of the ER by tandem interactions with hERG1a homotetramers. This model is reminiscent of the dimeric trafficking of sodium channel pairs (43). Alternatively, hERG1b may reside as homodimers that interact with newly synthesized hERG1a homodimers. This model is inspired by the “dimerization of dimers” mechanism previously established for homomeric assembly of Kv1.3 channels (44). When both hERG1a and 1b mRNAs are locally abundant, the ERSA may play less of a role as cotranslational heterodimers are formed, which then assemble into heterotetramers. Future work determining stoichiometry and composition under different conditions will be required to further illuminate the complexities of assembly.

Liquid-liquid phase separation (LLPS) serves to concentrate proteins and nucleic acids into condensates that regulate many cellular functions and play a critical role in physiology and disease (45, 46). Although most studies have focused predominantly on condensates of cytosolic proteins, growing evidence highlights phase separation of membrane proteins, including those involved in cell signaling (47-49). Our findings suggest that condensates may aid in preserving the integrity of heteromeric hERG channels, creating a protected environment that shields proteins from diffusible components of the proteasomal machinery. This environment, within ERSA, may help manage asymmetric hERG1a and 1b subunit production arising from uneven transcription initiated at their respective promoters (7). By serving as a reservoir, the condensate increases the local concentration of hERG1b, maximizing likelihood of successful association of these rare proteins in the crowded cellular environment.

Dynamics of the 1bNT fragment within condensates suggests this domain is capable of phase separation, but the full-length hERG1b ERSA lack liquid properties and may instead represent gel-like assemblies or arise from scaffolding by unidentified proteins that are involved in RXR-type ER retention. Thus, although hERG1bNT molecules can phase separate, we do not know whether this property is sufficient to drive condensation of the attached transmembrane domains in the ER. The tools available to test this hypothesis are limited (33), requiring future developments to determine whether phase separation drives the homotypic association of hERG1b membrane protein in the cellular environment.

Different sequence motifs regulate ER localization through either retention or retrieval mechanisms. Soluble proteins such as the ER chaperones protein disulfide isomerase (PDI) and binding immunoglobulin protein (BiP) can exit the ER but are later retrieved from the Golgi via COPI-coated vesicles. These vesicles contain KDEL receptors that recognize a KDEL motif harbored within the soluble proteins’ sequences (50, 51). Membrane proteins often use dibasic motifs such as KKXX, KXK and RXR to regulate ER localization. These motifs can mediate either retention within the ER, as observed for NMDA receptor subunits (52), or retrieval from the Golgi via COPI-coated vesicles, as reported for the GABA_B_ receptor and TREK2 K+ channel (53, 54). Our findings show that hERG1b puncta do not colocalize with COPI-positive compartments, suggesting that hERG1b sequestration by the N-terminal RXR motif does not involve Golgi-ER retrieval. We propose that the RXR helps establish the ERSA compartment to stably retain fully synthesized hERG1b proteins in the ER, shielding them from proteasomal degradation. Introducing an LQT2 mutation in hERG1b that misfolds the protein (25) precludes its localization to ERSA (and exposes it to the proteasome), suggesting that correct folding is a prerequisite for ERSA formation (Fig. 3 *E*). This novel mechanism suggests an expanded role for the RXR motif, linking classical quality control functions to biophysical compartmentalization to promote successful heteromeric assembly. It remains to be determined whether RXR-mediated ER retention promotes ERSA formation in other membrane proteins including the NMDA receptor and K_ATP_ channel subunits, where the RXR governs the ER exit of only fully realized oligomers (52, 55). Indeed, studies of such complexes could help establish a role for the ERSA compartment as fundamental to hetero-oligomeric assembly.

Why does hERG1b require an ER retention motif? We argue that hERG1b homomeric channels trafficking to the membrane of native cardiomyocytes could deleteriously affect cardiac I_Kr_. Cardiac I_Kr_ exerts its repolarizing effect late in the ventricular action potential (56). This critical delay is due to slow activation combined with rapid inactivation, suppressing currents during early depolarization. Later, as inactivation recovers, the current rebounds and facilitates repolarization (1, 2, 57). Homomeric hERG1b channels, however, activate rapidly and exhibit little inactivation, producing a classic delayed rectifier-type current (7). This current would counteract the depolarized plateau potential and shorten action potential duration (APD). Similar effects are observed with hERG agonists or mutants such as inactivation-impaired S631A, both of which are associated with short QT syndrome, in which the shortened APD allows reentrant arrhythmias in both the ventricle and atria (58-60). Although no pathogenic mutations in the hERG1b RXR signal have been reported to date, the essential role of hERG1 hetero-oligomerization underscores the importance of underlying mechanisms in maintaining cardiac rhythm.

## Methods

### Cell culture

HeLa cells (ATCC, CCL-2) were cultured in 35mm #1.5 glass bottom dishes (Cellvis, D35-14-1.5-N) using DMEM culture media (ThermoFisher, 11995065) supplemented with 10% FBS (ATCC, SCRR-30-2020). HeLa cells were transfected with 500 nanograms of hERG1a or 1b plasmid DNA per 35mm dish with 1 μl of TransIT-X2 Transfection reagent (Mirus Bio, MIR6004), and cells were imaged 48 h after transfection. Human iCell iPSC-CMs (CDI-FujiFilm) were thawed, plated, and cultured with CDI media according to the manufacturer’s specifications. After 10 to 14 days in culture, cells were transfected or fixed for immunostaining followed by imaging. For STED microscopy, mRNAs expressing SNAP tagged hERG1b and HaloTag containing KDEL motif in its C terminus were transfected to iPSC-CM using Lipofectamine MessengerMAX (LMRNA001). 100 nM SNAP-tag ligand JFX650 and 100 nM HaloTag ligand JFX549 were added to the culture media for 30 min, followed by three washes 5 min each using pre-warmed culture media to remove the free ligand. For membrane labeling, 100 nM Potomac Yellow dye was added to the cell culture media for 15 min, and the cells were imaged live.

### Plasmids and Generation of inducible stable cell line

Overexpression plasmids were made from pcDNA3.1 as a backbone. Truncations and insertion of coding sequences of hERG1a and 1b subunits and fluorescent proteins were performed using In-Fusion® Snap Assembly Master Mix (Takara, 638948). To generate a doxycycline-inducible cell line to express SNAP-tagged hERG 1b, we used a transcription activator-like effector nuclease (TALEN)-mediated targeting system, which incorporated the transgene at the AAVS1 locus (Qian et al., 2014). The puromycin-resistant donor cassette was used to express SNAP-tagged hERG1b, followed by an internal ribosome binding site and a gene encoding mApple. The cassette was transfected into HeLa cells with the two AAVS1-site specific TALEN plasmids at a ratio of 8:1:1, and cells were treated with puromycin (0.75 μg/ml) for approximately 3 weeks. Individual clones were picked and tested for doxycycline-induced expression of SNAP-tagged hERG1b and mApple. For overexpression studies, cells were treated with doxycycline (50 ng/ml) for 24-48 h prior to analysis.

### In vitro transcription of mRNAs

pCDNA3.1 plasmid containing SNAP-tagged hERG1b and Halo-KDEL with T7 promotor were linearized using EcoRI restriction enzyme at the 3’ end. Linearized plasmids were transcribed using mMESAGE mMACHINE™ T7 Transcription Kit (Invitrogen, AM1344) and poly(A) tailed using E. coli Poly(A) Polymerase (NEB, M0276S).

### Immunostaining

Cells were cultured in 12 mm coverslips with #1.5 thickness (Electron Microscopy Sciences 7229004). For labeling using HA (1:1000, Cell Signaling, 3724S), FLAG (1:1000, Sigma, F1804-1MG), Pan-hERG (1:200, Enzo Life Sciences, ALX-251-049-R100), and LC3A/B (1:400, Invitrogen, PA5-22731) antibodies, cells were fixed with 4% paraformaldehyde (PFA) followed by three 5-min washes using PBS. Cells were permeabilized using 0.5% Triton-X-100 followed by 30 min incubation in blocking solution (0.05% Tween-20 and 5% normal goat serum) and overnight incubation at 4°C with primary antibodies diluted in blocking solution, then incubation with secondary antibodies for 1 h at room temperature. For labeling using Sec24a (1:300, Santa Cruz Biotechnology, sc-169279), COPB (1:300, Santa Cruz Biotechnology sc-393615) and LAMP1(1:100, Invitrogen, eBioH4A3) antibodies, cells were fixed using 100% ice-cold methanol for 10 min at -20°C, followed by three 5-min washes in PBS and blocking and incubation with primary antibody overnight at 4°C, then secondary antibody for 1 h at room temperature.

### Microscopy and image analysis

Live and fixed confocal imaging was performed using a Nikon Ti2 spinning disk confocal microscope with a 60x (1.4NA) and 100x (1.45NA) oil immersion objective and Hamamatsu ORCA-Flash4.0 sCMOS camera. Nikon Elements software was used for image acquisition.

Puncta quantification and colocalization analysis were performed using the Imaris (Oxford instruments) spots model. Spot detection was initialized using an average estimated diameter of 500 nm and modeled PSF-elongation along the z-axis with an estimated diameter of 1.2 μm. Initial filtering was applied using the *quality* parameter with an empirically determined threshold to maximize the inclusion of true-positive puncta while excluding background noise. Additionally, spot regions were thresholded based on *local contrast* with an empirically determined threshold to refine spot regions. To ensure consistency and reproducibility, configuration files containing all the thresholding parameters were generated and applied across images, enabling unbiased segmentation. Further, the spots were filtered for those greater than 300 nm in diameter in the XY plane. Puncta were detected throughout the field of view of the microscopy images and assigned within individual cellular boundaries to quantify puncta on a per-cell basis.

For 1bNT condensate experiments, the number of cells with condensates was quantified manually in a blinded manner. All the quantifications were conducted on raw images, and sample images provided in the figures were deconvoluted using Huygens professional software (Scientific Volume Imaging), using the Classic Maximum Likelihood Estimation (CMLE) algorithm with default settings. Deconvolution was applied evenly to the images used for comparison.

Leica TCS SP8 STED system equipped with highly sensitive HyD detectors and an HC Plan Apo CS2 100X, 1.4NA oil immersion objective lens was used to acquire STED images. Random regions of interest (25 x 25 μm squares) were selected for image acquisition in each cell. Acquisition parameters were controlled by the Leica Application Suite platform (LASX) and images were deconvolved using Huygens CMLE algorithm. Puncta with ER colocalization analysis was performed using the Imaris spots and surface model. The number of puncta colocalized within the 50 nm, 100 nm, or 200 nm threshold distance from the center of the puncta to the surface was quantified using Imaris.

Fluorescence recovery after photobleaching (FRAP) studies were performed using a Nikon Ti2 spinning disk confocal equipped with a 37° C incubator and 5% CO_2_ supply. 100 μs stimulation with maximum power using a 488 nm stimulation laser was used to photobleach the region of interest and images were acquired before and after photobleaching in 2-sec intervals for 3-5 min. ImageJ was used to measure the intensity of the photobleaching area and plotted against time.

### Pulse-chase analysis of hERG1 subunits

In the pulse-chase analysis, we transfected plasmids containing either hERG1a-SNAP-IRES-Halo, hERG1b-SNAP-IRES-Halo, hERG1a-Y611H-SNAP-IRES-Halo or hERG1b-Y271H-SNAP-IRES-Halo. 24 h after transfection, the proteins were labeled with 100 nM JFX650 SNAP-Tag ligand and 100 nM JF479 HaloTag ligand, followed by three 10-min washes. Subsequently, 100 nM PA-JF549 SNAP-Tag ligand was added and left in the media until the end of the experiment. The live cells were imaged immediately (0 h) and at 2-h intervals up to 8 h after labeling using a Nikon Ti2 spinning disk confocal microscope equipped with a 37° C incubator and 5% CO_2_ supply. The HaloTag expression was used to identify transfected cells even after complete degradation of hERG proteins. ImageJ was used to manually segment the cells, and the integrated density values were exported per cell. Corrected total cell fluorescence (CTCF) was calculated using the formula: CTCF = Integrated Density – (Area of selected cell X Mean fluorescence of background). Mean CTCF values were normalized by setting the baseline mean CTCF at 0 h to 100 for plotting one phase exponential decay.

### Statistical analysis

Statistical tests were performed using GraphPad Prism software (Dotmatics) with unpaired Student’s t-tests for comparing two groups and one-way ANOVA tests for comparing multiple groups. Statistical tests for each data set are listed in the figure legends.

## Acknowledgments

We thank members of the Robertson lab, especially Catherine Eichel, Lisandra Flores-Aldama, and Taylor Voelker for critical discussion, Fang Liu for technical support, and members of the Audhya lab for helpful advice and reagents. Zachary T. Campbell and Cynthia M. Czajkowski for reviewing the manuscript and providing suggestions. We thank Luke D. Lavis for providing Janelia Fluor dyes. The work was supported by NIH grants R01-HL131403 and R01-NS081320 (G.A.R.), American Heart Association Predoctoral Fellowship 24PRE1188958 (S.K.) and NIH grants R01 NS124165 and R35 GM134865 (A.A).

