## Supplemental figures for "A privileged ER compartment for post-translational heteromeric assembly of an ion channel"

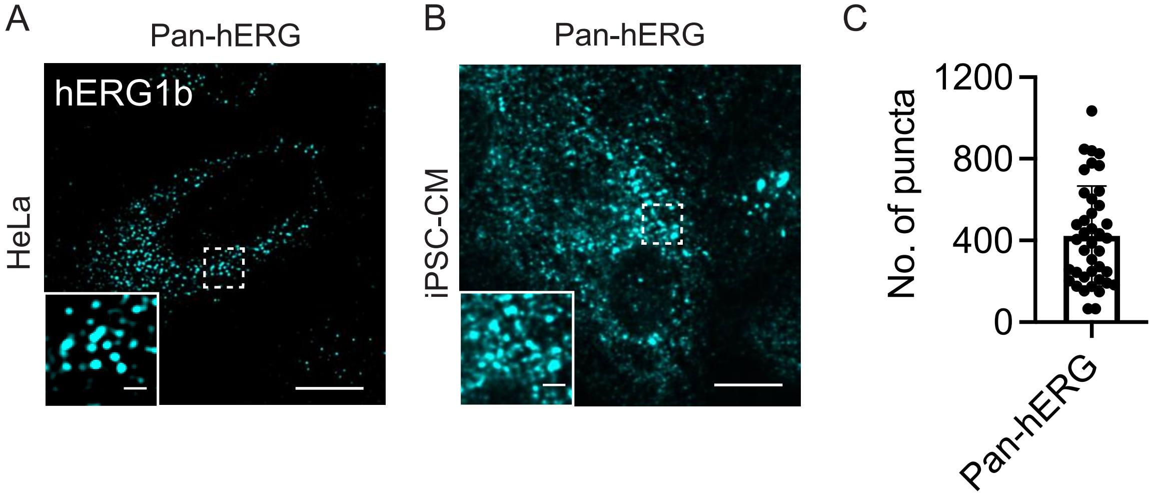


**Fig. S1.** Untagged hERG channels form puncta. (*A*) HeLa cells transfected with untagged *hERG1b* plasmid and stained with pan-hERG antibody; (*B*) confocal image of endogenous hERG1a and 1b in fixed iPSC-CM stained with pan-hERG; (*C*) quantification of number of puncta per iPSC-CM (n= 40 cells) from one representative experiment, repeated twice; scale bar: 10 μm in large image and 2 μm in inset.

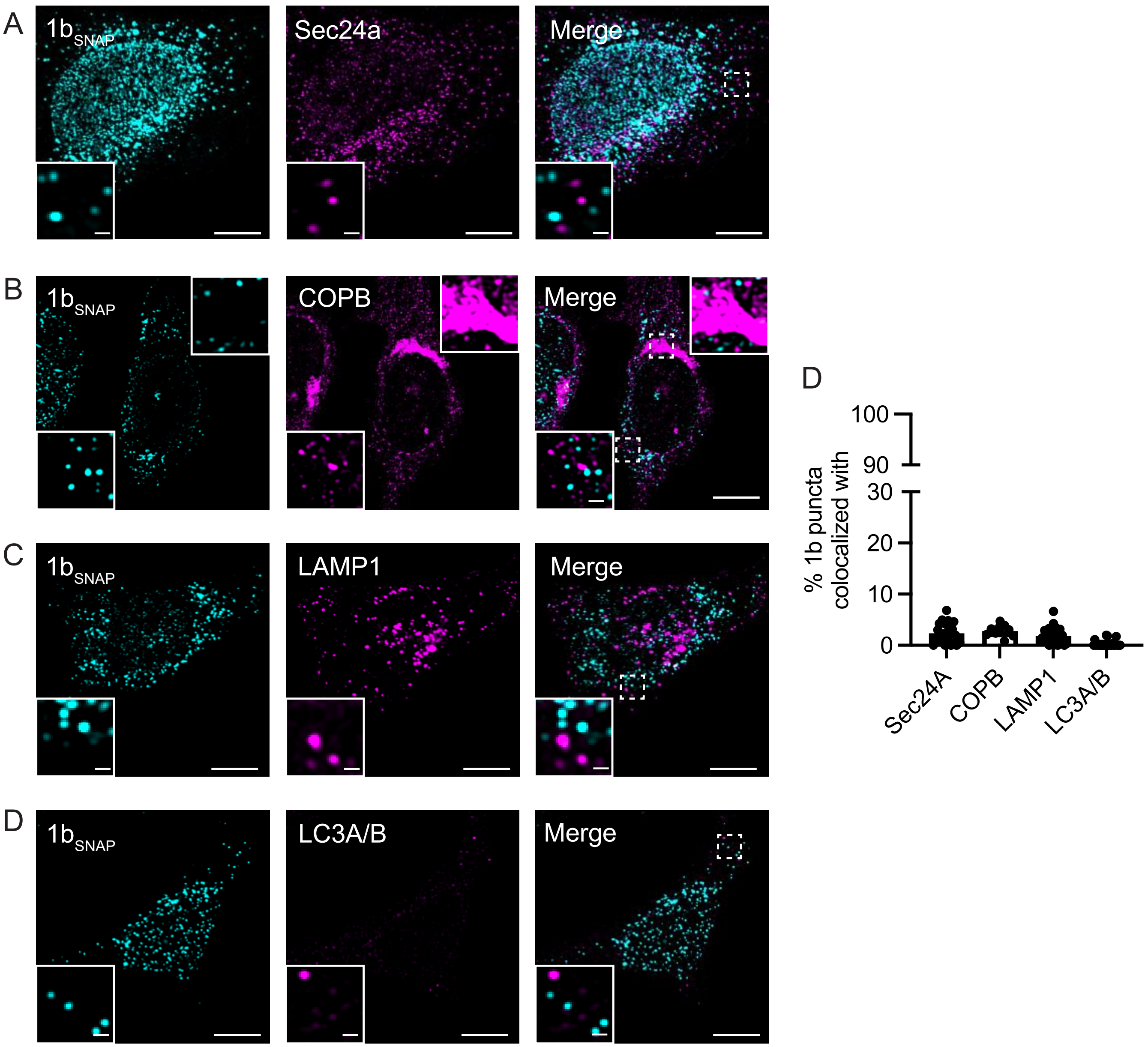


**Fig. S2.** hERG1b puncta do not colocalize with other subcellular compartments. Confocal images of a stable HeLa cell line expressing hERG1b$_{SNAP}$ co-labeled with (*A*) Sec24a antibody labeling COPII (ER-Golgi) vesicle (1), (*B*) COPB antibody labeling COPI (Golgi-ER) vesicle (2), (*C*) LAMP1 antibody labeling lysosomes (3), (*D*) LC3A/B antibody labeling autophagosomes (4); (*E*) percentage of hERG1b puncta colocalized with each marker; colocalization analysis was carried out by thresholding the distance between the centroids of hERG1b puncta and other compartment at 250 nm; percentage of hERG1b puncta colocalized within 250 nm of the ER marker Halo-KDEL in iPSC-CM is also given for comparison. Data are mean ± s.d. (n=24 cells per condition); Scale bar: 10 μm in the large image and 1 μm in the inset. hERG1b puncta rarely colocalized with COPII, COPI (both tubular and vesicle fractions), lysosomes or autophagosomes indicate that hERG1b puncta are exclusive to ER. Indeed, the cells did not have many autophagosomes, indicating hERG1b expression did not trigger cellular stress, at least not within 48 h after inducing the expression (5).

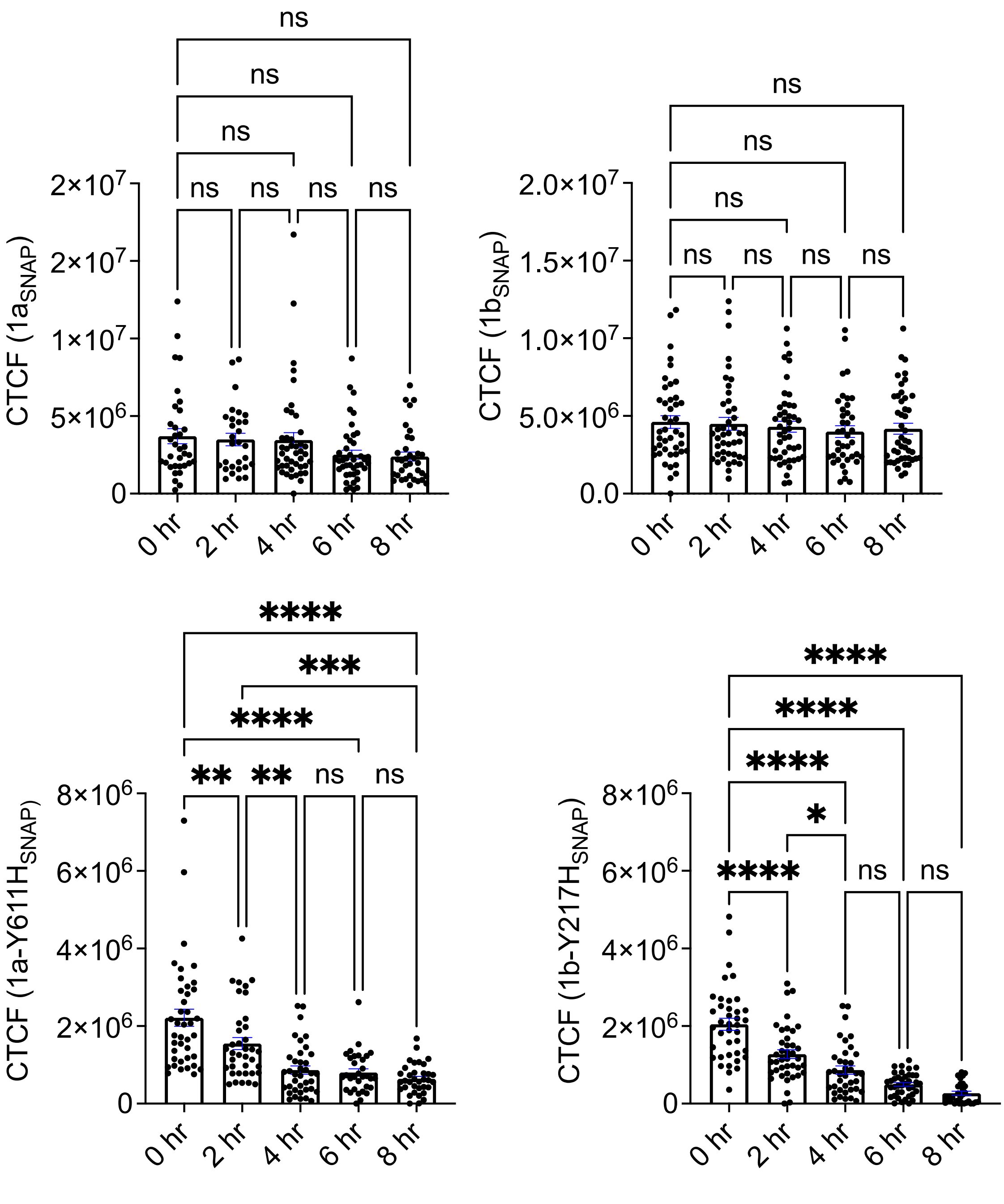


**Fig. S3.** hERG1b is sequestered in a privileged compartment. Corrected total cell fluorescence intensity (CTCF) quantified across time from HeLa cells transfected with either (*A*) hERG1a$_{SNAP}$, (*B*) 1b$_{SNAP}$, (*C*) 1a-Y611H$_{SNAP}$, or (*D*) 1b-Y271H$_{SNAP}$; data are mean ± S.E.M, n= 35-45 cells per condition; analyzed with a one-way ANOVA (**** indicating $p<0.0001$, * indicating $p<0.05$, ns, not significant).

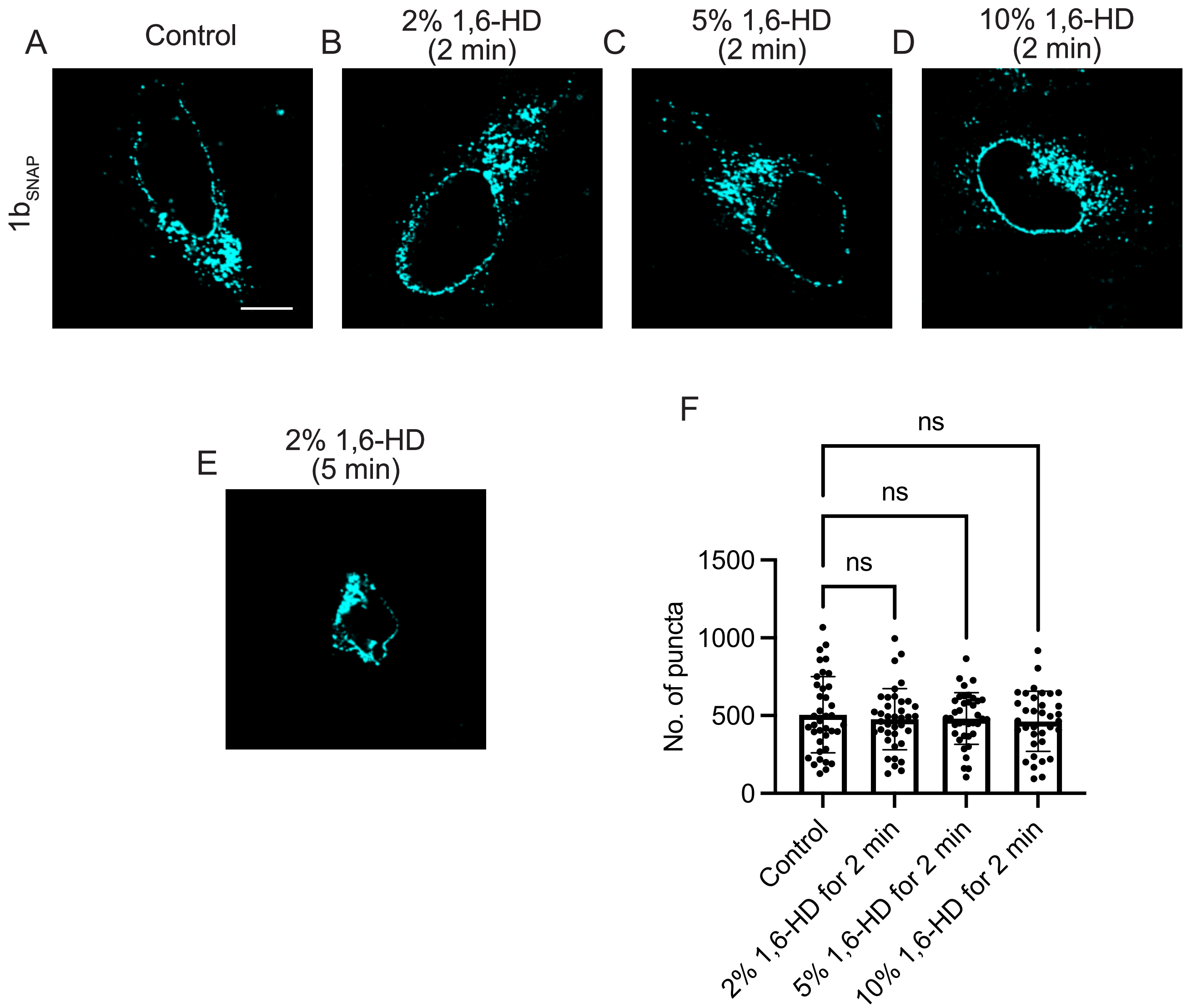


**Fig. S4.** 1b puncta are unaffected by 1,6-hexanediol. Confocal images of HeLa cells expressing $1b_{SNAP}$ incubated with 1,6-hexanediol per following conditions: *A*) Control; B) 2%, 2 min; C) 5%, 2 min; *D*) 10%, 2 min; *E*) 2%, 5 min. (*F*) quantification of number of puncta per cell; data are mean ± s.d., n=35-38 cells per condition, analyzed with one-way ANOVA (ns indicating no significance); scale bar: 10 µm. 1,6-Hexanediol was directly dissolved in the DMEM media w/FBS, pre-warmed, added to the cells, and incubated for the indicated time. After incubation, the cells were fixed with 4% PFA for 5 min and washed 3 times with 1x PBS before imaging. Typical concentrations (2-10%) used in the literature (6-8) did not disrupt hERG1b puncta within 2 minutes of incubation. More prolonged incubation (5 min) at a 2% concentration affected cellular morphology in HeLa cells, as previously reported (6). This result indicates that the 1b puncta may arise from interactions other than weak hydrophobic interactions (9).
